# EssTFNet: Integration of Adaptive Time–Frequency and DNA Language Models for Interpretable Human Essential Gene Prediction

**DOI:** 10.64898/2025.12.31.697261

**Authors:** Dong-Xin Ye, Shi-Shi Yuan, Wei Su, Hong-Qi Zhang, Rui Li, Ye-Chen Qi, Nan-Qing Dong, Hao Lin, Yan-Ting Jin

## Abstract

Essential genes are defined as those that are indispensable for an organism’s survival. The loss of function of these genes results in cell death or an inability to complete the normal life cycle. Research on essential genes is pivotal in elucidating the origin and evolution of life, as well as in identifying potential therapeutic targets. Therefore, accurately predicting essential genes is of great scientific importance and has many applications in basic research and the biomedical field. In this study, we propose EssTFNet, a novel and interpretable deep learning framework that combines adaptive time–frequency analysis with a DNA language model to achieve accurate prediction of human essential genes while enabling mechanistic biological interpretation. Specifically, EssTFNet leverages the architecture of ATFNet, which innovatively maps DNA and protein sequences into equivalent time-series signals to extract periodic and non-stationary features, thereby enhancing the model’s capacity to capture complex sequence patterns. Through effective feature selection and architectural optimization, EssTFNet strikes a favorable balance among prediction accuracy, model interpretability, and cross-tissues generalization capability, significantly outperforming current mainstream sequence-based deep learning methods with an AUC of 0.8336 and an AUPR of 0.8212. Additionally, the DeepLIFT attribution method was employed to identify functional motifs closely associated with gene essentiality, offering valuable insights for experimental validation. For the convenience of researchers, we have developed an easy-to-use web server and made it along with the source code in a GitHub repository: https://github.com/QIANJINYDX/EssTFNet. Overall, this study presents a potentially useful methodological framework for human essential genes prediction, which could provide valuable insights for future research and applications in this field.

## 1 Introduction

Essential genes are indispensable to the survival or reproduction of a living organism, playing a crucial role in maintaining cellular life. Studying these genes can help to elucidate the origin and evolution of life, define the minimal gene set required for cellular viability, and infer the genomic architecture and lifestyle of early life forms (Zhang et al. 2016). Moreover, identifying essential genes in pathogens is key to drug design, as antibiotics primarily target the fundamental metabolic processes of pathogens, making essential genes potential drug targets (Hutchison et al. 2016; Qiao et al. 2024; Qi et al. 2025). Similarly, identifying essential genes in cancer cells can aid in discovering novel cancer drug targets (Hu et al. 2007; Wei et al. 2014; Wang et al. 2015). Therefore, the identification of essential genes is substantial scientific significance and broad application value in both basic research and biomedical fields.

Currently, there are two main approaches for identifying essential genes: experimental and computational methods. Experimental methods can provide specific results for essential genes under different experimental conditions (Liang et al. 2024), such as RNA interference technology, CRISPR-based genome editing, gene knockout, transposon mutagenesis, and others (Giaever et al. 2002; Salama et al. 2004; Chen et al. 2015; Ma et al. 2015). Although highly reliable, these methods are labor-intensive, costly, and technically demanding, especially in complex organisms (Li et al. 2020).

To overcome these limitations, computational methods have been developed to predict gene essentiality using machine learning algorithms (Wang et al. 2024; Xie et al. 2025) and biological feature data(Mohapatra and Sahu 2025; Zeng et al. 2025). This approach significantly reduces the experimental burden while accelerating the discovery process. For example, in 2005, Chen et al. studied the dispensability of proteins in Saccharomyces cerevisiae by combining high-throughput data and machine learning methods (Chen and Xu 2005). In 2006, Seringhaus et al. reported a machine learning-based method that integrates various intrinsic and predicted features to identify essential genes in the Saccharomyces cerevisiae genome (Seringhaus et al. 2006). In 2012, Yuan et al. integrated information-rich genomic features and used three machine learning-based methods to predict lethal knockouts in mice (Yuan et al. 2012). In 2015, Lloyd et al. analyzed the characteristics of essential genes in the Arabidopsis thaliana genome and used Arabidopsis as a machine learning-based model to transfer essential annotations to rice and Saccharomyces cerevisiae (Lloyd et al. 2015). In 2017, Guo et al. proposed a method that accurately predicts human essential genes using only nucleotide composition and associated information (Guo et al. 2017). In 2019, Tulio L. Campos et al. performed large-scale annotation and exploration of eukaryotic genes and protein functions, predicting essential genes both within and across species using machine learning methods (Campos et al. 2019). In 2020, Zhang et al. proposed the DeepHE method, which accurately predicts human essential genes by integrating sequence data and PPI network features, significantly outperforming traditional machine learning models (Zhang et al. 2020). In 2023, Li et al. proposed the DeepCellEss framework, which combines convolutional neural networks and self-attention mechanisms to provide interpretable deep learning methods specific to cell lines (Li et al. 2023).

While these studies highlight the feasibility of predicting essential genes computationally, several key challenges remain for human essential gene prediction: (1) Compared to microbial genomes, the human genome contains much longer and structurally complex sequences, posing significant difficulties for computational modeling (Lander et al. 2001). Therefore, it is essential to develop an efficient and accurate long-sequence modeling approach. (2) Although several DNA language models have been developed (Luo et al. 2025), there is no model specifically designed for essential genes. Therefore, building a language model focused on essential genes is of great significance. (3) Current research on human essential gene prediction still lacks in-depth mechanistic analysis, leading to limitations in prediction accuracy and interpretability. How to mine and extract biologically meaningful key features in computational analysis, and improve the algorithm’s ability to understand and utilize data features, is an urgent issue that needs to be addressed. (4) Most existing gene essentiality prediction models use a binary classification approach, distinguishing genes as either “essential” or “non-essential”. However, in certain task scenarios (e.g., drug target gene screening), gene essentiality often exhibits a continuous variation rather than a simple binary state (Hogan and Cardona 2022; Clyde 2023). Models capable of predicting the degree of essentiality would provide more nuanced insights, enhancing their applicability to drug discovery and personalized medicine (Chen et al. 2025).

To address these challenges, we propose EssTFNet, an interpretable hybrid framework that fuses feature representations from the adaptive time–frequency model and DNA language model to improve the prediction of human essential genes and their fitness scores. The overall research workflow of our proposed method is illustrated in Figure 1. By combining sequence-context embeddings with time–frequency features, EssTFNet improves predictive performance and provides additional clues for interpreting which sequence patterns contribute to the predictions. In our experiments, EssTFNet consistently outperforms existing methods for human essential-gene prediction. Overall, this work provides a practical computational tool for essential-gene prioritization and a mechanistically grounded basis for downstream applications in precision medicine, drug-target discovery, and synthetic biology.

**Figure 1.**
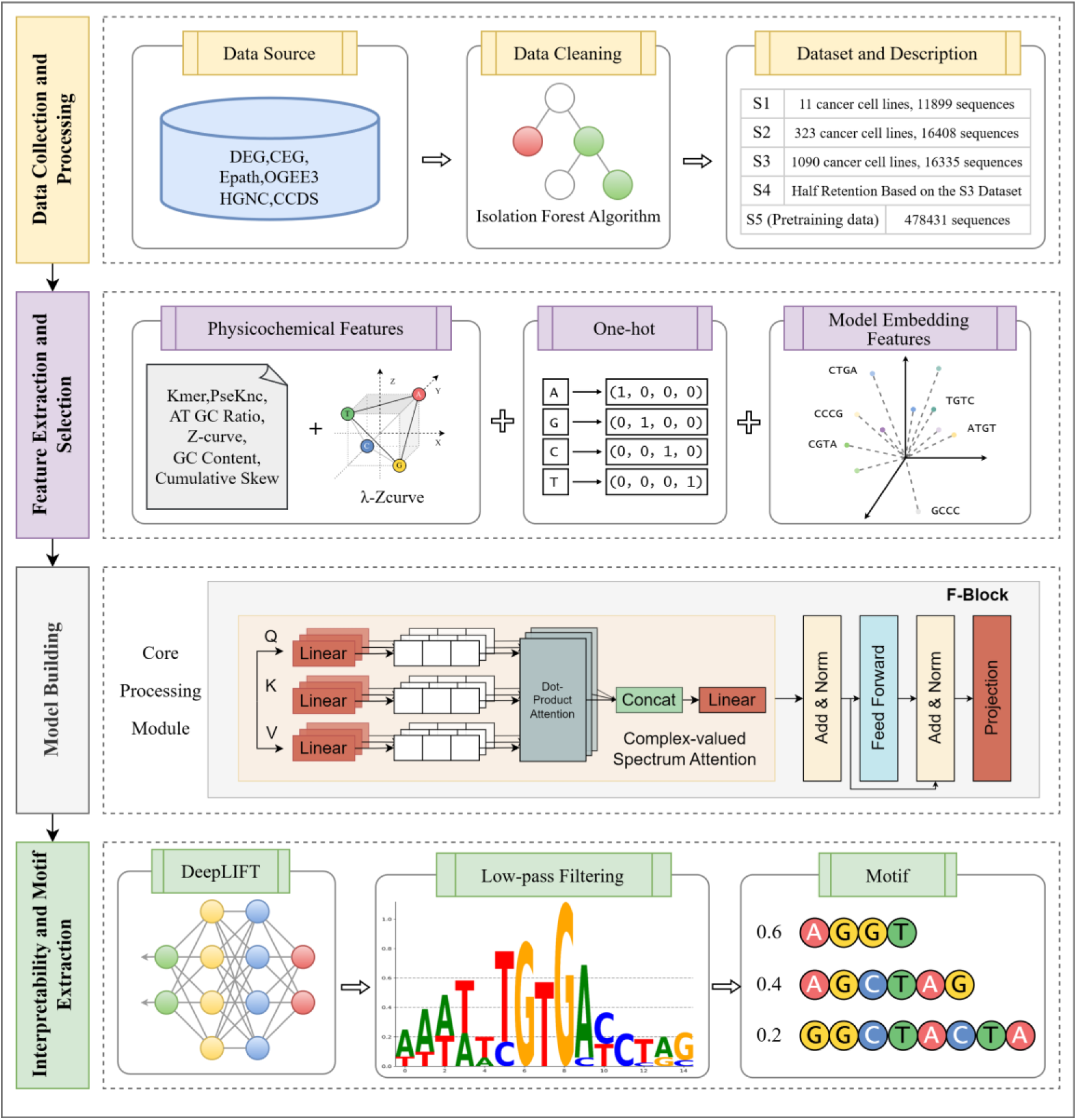
Overview of the EssTFNet workflow. Data collection and processing: Publicly available essential-gene resources (e.g., DEG/CEG, OGEE, EPath, OGEE3, HGNC, and CCDS) are collected and curated to construct the study datasets, followed by data cleaning and quality control. Feature extraction and selection: Three complementary feature streams are generated, including physicochemical descriptors (e.g., k-mer/PseKNC, AT/GC ratio, Z-curve, GC content, cumulative skew, and λ-Z-curve), nucleotide one-hot encoding, and embeddings extracted from a DNA language model. Model building: These features are integrated within EssTFNet, in which stacked F-blocks serve as the core building units, and additional modules are employed to perform multi-source feature fusion and final prediction. Interpretability and Motif Extraction: DeepLIFT attribution, low-pass filtering, and motif extraction are used to highlight sequence patterns associated with model decisions.

## 2 Results

### 2.1 Evaluation of EssTFNet

Our model achieved a significant performance improvement on the S1 dataset with an AUC of 93.27%. This represents a 4.73 percentage point increase over the method proposed by Guo et al., which reported an AUC of 88.54% (Guo et al. 2017). This result not only demonstrates our model’s superior discriminative capability but also visually showcases the stable superiority of our approach across varying feature quantities through the AUC comparison in Figure 2A.

**Figure 2.**
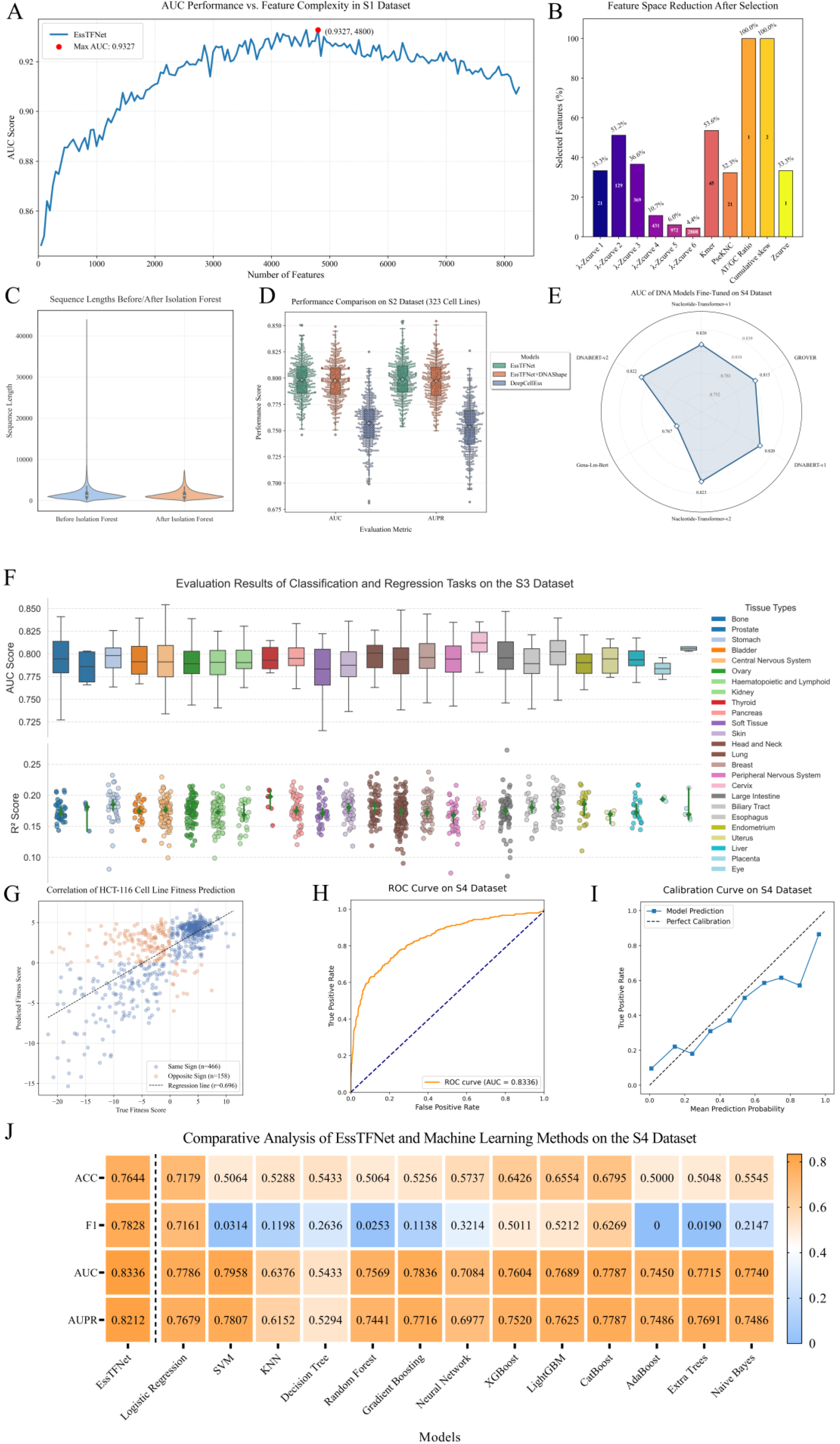
Comprehensive Evaluation of EssTFNet’s Performance Across Diverse Prediction Tasks. **A**: A curve illustrates the relationship between feature complexity and model performance for the EssTFNet algorithm on the S1 dataset. **B**: Isolation Forest outlier removal drastically reduces raw sequence lengths from over 40,000 bp (pre-processing) to near-zero values (post-processing), as shown by paired bar plots. **C**: Stacked bars quantify post-selection retention rates across feature categories. **D.** Compares the performance distribution of three models on the S2 dataset. **E.** Benchmarking results of different DNA language models. **F.** Displays model evaluation results for classification and regression tasks across multiple tissue types on the S3 dataset. The upper section shows box plots of AUC scores, while the lower section presents scatter plots of R² scores, with distinct colors distinguishing tissue types. **G.** Visualization of regression task predictions on the HCT-116 cell line. sequence-level scatter (n=624) with color indicating sign consistency—same sign (prediction and truth share the sign; blue) versus opposite sign (signs differ; orange). **H.** ROC curve on the S4 dataset. **I.** Calibration curve on the S4 dataset. **J.** Comparison of the model with different machine learning algorithms.

A detailed decomposition of this optimal feature subset is presented in Figure 2B. Notably, λ-Z-curve features dominate the selected subset, accounting for 4,730 of the 4,800 features, highlighting their critical role in the model’s decision-making process. Furthermore, to ensure modeling stability, we addressed the issue of anomalously long sequences within the raw S1 dataset. As shown in Figure 2C, we employed the Isolation Forest algorithm to identify and filter out these outliers, resulting in a more uniform and robust distribution of sequence lengths.

On the more challenging S2 dataset, our model demonstrated significant advantages, achieving an overall performance improvement of approximately 5% in both the AUC (from 75.69% to 79.82%) and AUPR (from 75.32% to 79.88%) compared to the DeepCellEss method, as shown in Figure 2D. These results demonstrate the robustness and generalizability of our model when handling complex biological data. To further examine feature sensitivity, we evaluated the impact of incorporating DNASHAPE structural features. A slight performance decrease of 0.08% (79.74%) was observed (Li et al. 2024) , which suggests two potential factors: (1) DNASHAPE features may partially overlap with the existing sequence-based features, introducing redundancy; (2) the current network architecture has room for improvement in its ability to integrate higher-order structural features. These findings provide important directions for future research, including refinement of feature selection strategies and optimization of heterogeneous feature fusion mechanisms.

We evaluated various existing DNA language models on the S4 general essential gene dataset, including DNABERT, DNABERT-2, GROVER, Nucleotide Transformer-v1, NT v2, and Gena-LM-BERT. Among these, NT v2 demonstrated the best performance in terms of the AUC metric and was therefore selected as the base model (see Figure 2E for comparison results).

Our model also demonstrated strong adaptability on the larger-scale and more organizationally diverse S3 dataset, which included 1,090 cell lines and 25 tissue types (Figure 2F). It achieved an average AUC of 79.29% in classification tasks. Although performance declined by only 0.53% compared with the S2 dataset (323 cell lines), the model maintained robust generalization capabilities, indicating its stability across data of varying scales and tissue origins. Complementing these classification results, we next evaluated regression performance for predicting continuous fitness values. Although the overall R² suggests room for improvement in absolute accuracy, the model captured trends well on the HCT-116 cell line: 74.6% of genes had correctly predicted signs, and the Pearson correlation reached 0.696. Moreover, the model demonstrated superior predictive performance for essential genes, which clustered in the upper-right region (high Fitness values) of Figure 2G, indicating its outstanding capability in identifying functionally critical genes. These findings highlight both the current limitations of regression predictions and the practical potential of our approach for functional genomic applications.

The ROC curve presents the performance of the general essential gene prediction model (S4) on the test set. The model attains an AUC of 0.8336, demonstrating strong discrimination between essential and non-essential genes (Figure 2H). This exceeds the performance of cell line–specific S3 models, highlighting the robustness and generalizability of the unified S4 model across diverse contexts. The high true positive rate across a broad range of false positive rates further supports the effectiveness of the model in general essential gene identification. Figure 2I displays the calibration curve of the model on the S4 dataset, assessing the agreement between predicted probabilities and the actual occurrence of positive class instances. Overall, the prediction curve closely follows the ideal calibration line, indicating reliable probabilistic outputs and providing robust support for gene functional studies.

On the S4 dataset, this study conducted a comprehensive comparative analysis between the proposed EssTFNet model and 13 widely used traditional machine learning methods using multiple evaluation metrics. As illustrated in Figure 2J, EssTFNet demonstrated significant advantages across all four key indicators: it achieved an accuracy of 0.7644, an F1 score of 0.7828, an AUC of 0.8336, and an AUPR of 0.8212—all substantially outperforming the competing methods. In contrast, most traditional models showed clear performance deficiencies on this dataset, particularly in terms of the F1 score, which is especially sensitive to class imbalance. Statistical results indicate that up to nine traditional models recorded F1 scores below 0.5, with some approaching zero, highlighting notable performance bottlenecks in addressing this task. Although a few traditional models performed relatively well in AUC and AUPR metrics, they still fell short when evaluated across all dimensions. These findings further emphasize EssTFNet’s superior capability in feature extraction and classification for complex and imbalanced time-series data, showcasing its comprehensive performance advantages.

In summary, these experimental findings comprehensively validate EssTFNet’s robust predictive capability derived from its deep feature fusion and spatiotemporal modeling architecture, particularly its exceptional competence in capturing intricate biological feature relationships through combined temporal-frequency domain analysis.

### 2.2 Model interpretability analysis

We performed t-SNE dimensionality reduction projection and clustering analysis on various features before model fusion to evaluate the performance of different feature types in distinguishing essential genes from non-essential genes, as shown in Figure 3A-C. The results show that the DNA physicochemical property features exhibit a clear clustering trend in the two-dimensional space, which can distinguish the two types of gene samples to some extent. This likely reflects intrinsic differences in structural stability, base composition, and other biophysical properties between these two gene categories. In contrast, protein-level features exhibited relatively scattered distributions, with no clear separation observed. After feature integration via model fusion, a comprehensive t-SNE analysis revealed clear category boundaries in the low-dimensional space, confirming the effectiveness of multi-source heterogeneous feature fusion in enhancing discriminative capability. This further indicates that the deep features extracted by the model have a strong capacity to retain class discriminative information, providing a solid foundation for downstream prediction and functional interpretation.

**Figure 3.**
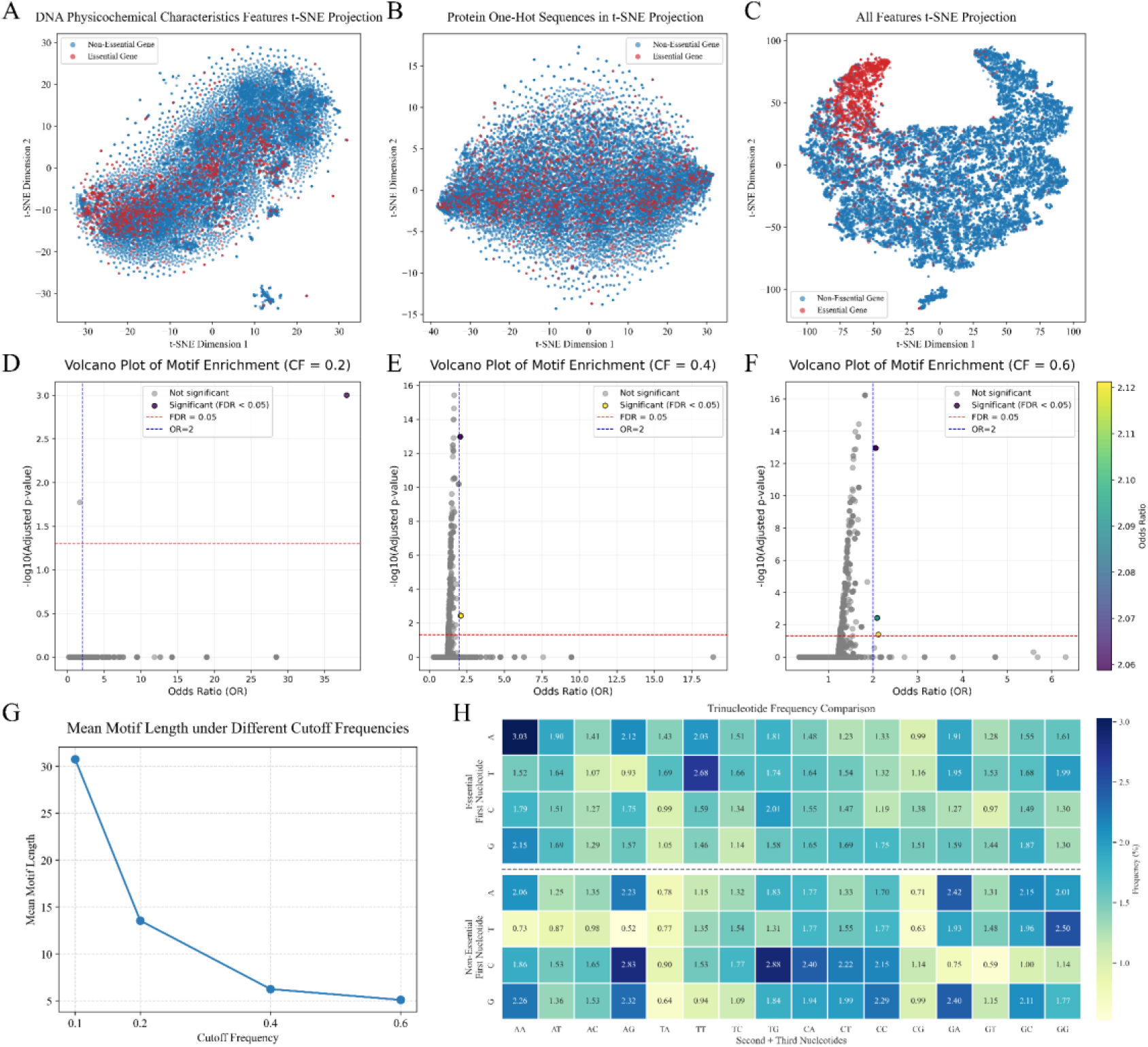
Analyzing and interpreting the model. **A-C**. t-SNE embeddings of feature spaces: (A) DNA physicochemical features, (B) protein one-hot sequences, and (C) their combination for essential-gene prediction; each point is a sequence (red = essential, blue = non-essential).**D-F**. Volcano plots of motif enrichment under dual-validation (Fisher’s exact test + Benjamini-Hochberg) **G**. Increasing the Butterworth low-pass cutoff from 0.1 to 0.6 sharply reduces the mean motif length (from >30 bp to ∼5 bp), demonstrating a tunable mechanism for extracting motifs of different sizes. **H**. Trinucleotide frequency comparison between essential and non-essential genes.

The identified motifs were filtered using a two-step filtering strategy. Results were visualized through volcano plots, where the y-axis represents -log10 (adjusted p-value) and the x-axis shows OR values on a log2 scale, with reference lines marking FDR thresholds and effect size cutoffs (Figure 3D-F). Significantly enriched motifs clustered in the upper-right quadrant denote high-confidence positive regulatory elements.

Figure 3G illustrates the relationship between the cutoff frequency and the mean motif length extracted using the Butterworth low-pass filter. As the cutoff frequency increases from 0.1 to 0.6, the mean motif length decreases significantly—from over 30 bp at a cutoff of 0.1 to approximately 5 bp at a cutoff of 0.6. This trend confirms the effectiveness of the low-pass filtering approach: lower cutoff frequencies allow the extraction of longer motifs by preserving low-frequency signals and filtering out high-frequency noise. Conversely, higher cutoff frequencies retain more high-frequency components, which results in the identification of shorter motifs. These results demonstrate that adjusting the cutoff frequency provides a tunable mechanism for targeting motifs of varying lengths, supporting the dynamic configuration strategy based on expected motif size.

To explore the differences in nucleotide sequence composition between essential and non-essential genes, we statistically analyzed the frequency of all 3-mer (trinucleotide) combinations in both groups, as shown in Figure 3H. Non-essential genes were sourced from the S4 dataset, while essential genes were integrated from the S5 dataset. The analysis revealed that the 3-mer sequence “AAA” occurred significantly more frequently in essential genes than in non-essential genes. Notably, the 3-mer “AAA” encodes the amino acid lysine (Lys) during the post-transcriptional translation process. Lysine is a basic essential amino acid that cannot be synthesized from other metabolic intermediates in mammals and must be obtained through external intake (Tomé and Bos 2007). Lysine often resides in functional regions of protein structures, such as nuclear localization signals (NLS) or DNA/RNA binding sites, and is involved in regulating various cellular processes like chromatin remodeling and transcriptional regulation (Chen et al. 2022). This enrichment phenomenon may reflect the high dependency of essential genes on lysine residues in terms of functionality, suggesting their critical role in maintaining basic life functions. Additionally, the motif recognition results in Figure 4 also show that many high-frequency motifs are rich in adenine (A), further supporting the above observations. This finding validates the biological reasonableness and reliability of the motifs identified by the model.

**Figure 4.**
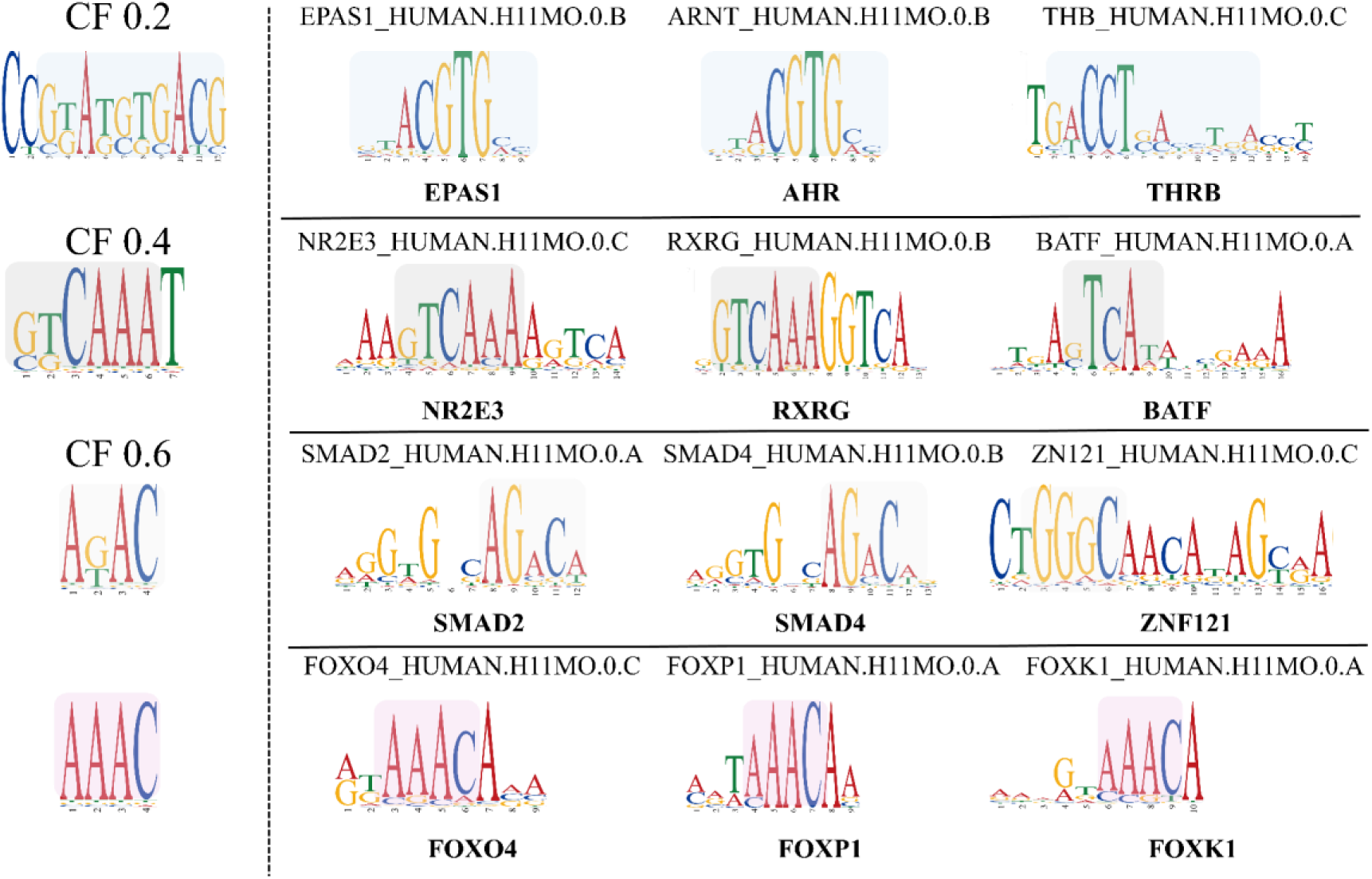
This chart displays multiple human genes and their corresponding motif segments. On the left are essential motifs identified at different CF (cutoff frequency) thresholds, while the right side shows highly similar motif fragments along with their corresponding genes.

### 2.3 Motif analysis

In addition, we performed motif discovery and interpretability analysis. Following multiple rounds of parameter optimization and motif screening, we ultimately identified four non-redundant DNA motifs by merging redundant items, including two long motifs and two short motifs, as shown in Figure 4. Subsequently, we performed motif similarity comparison using the Tomtom online platform, selecting all human DNA sequences for alignment against the HOCOMOCO Human database(Gupta et al. 2007; Kulakovskiy et al. 2018). The top three matches with the highest similarity for each motif were selected for analysis, with the corresponding related genes shown on the right side of Figure 4. The long motifs typically contain more complex structures and multiple DNA-binding domains, enabling them to regulate the expression of multiple genes. (Das and Dai 2007). For instance, EPAS1 (associated with hypoxia response and angiogenesis) may influence cellular survival and environmental adaptation capabilities by regulating oxygen-sensing mechanisms and angiogenic pathways(Beall et al. 2010).AHR, a core regulatory node linking immunity, metabolism, and oncogenesis, orchestrates essential gene expression programs through stress response modulation and metabolic rewiring, thereby enabling cells to dynamically adapt to microenvironmental challenges(Ema et al. 1994; MacPherson et al. 2013; Gomez et al. 2018; Sadik et al. 2020). Long-motif transcription factors typically mediate indispensable functions for maintaining core cellular physiology. They often exhibit direct interactions with essential genes to sustain basal survival and metabolic functions. In contrast, short motifs, composed of a limited number of amino acids in their DNA-binding or transcription factor-binding domains, generally demonstrate stringent specificity and precisely defined target loci. For example, SMAD2 and SMAD4, key components of the TGF-β signaling pathway, regulate cell proliferation, differentiation, and apoptosis(Eppert et al. 1996; Yang et al. 2015). Dysregulation of this pathway is frequently implicated in cancer and other pathologies. Within the essential gene network, these genes may act as pivotal nodes modulating the equilibrium between cellular survival and demise. By integrating gene regulatory information from both long and short motifs, the model delivers more comprehensive and precise analytical outcomes, enabling in-depth exploration of the multilayered roles genes play in cellular physiological processes. This approach holds significant potential for advancing the understanding of gene regulatory networks, investigating disease mechanisms, and driving innovations in personalized medicine.

### 2.4 Ablation study

To investigate the contributions of various components within EssTFNet, we conducted a comprehensive ablation study, as summarized in Table 1. The complete EssTFNet model achieved the highest performance across all evaluation metrics, with an Accuracy of 0.7644, F1-Score of 0.7828, AUC of 0.8336, and AUPR of 0.8212, thereby demonstrating the effectiveness of its fully integrated design.

**Table 1.**
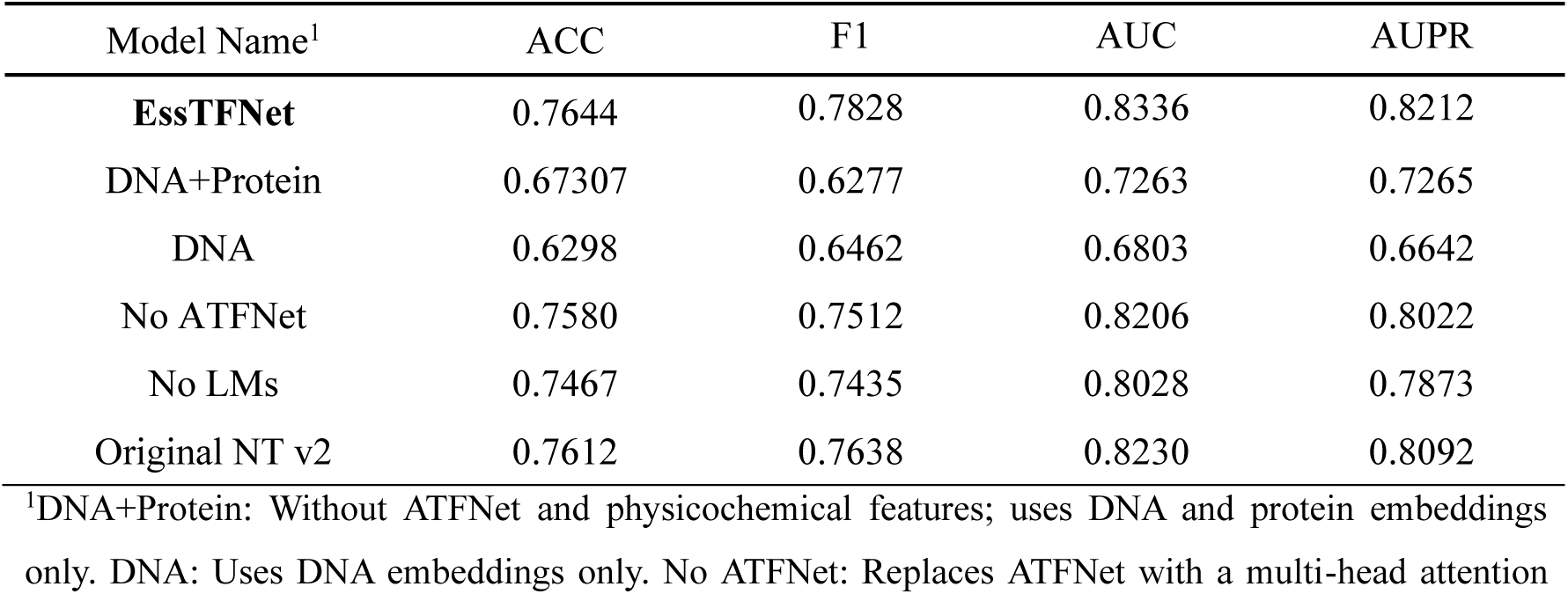
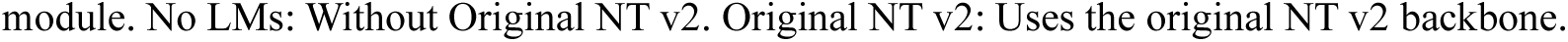
Ablation Study of EssTFNet Modules on Prediction Performance.

| Model Name<sup>1</sup> | ACC | F1 | AUC | AUPR |
| --- | --- | --- | --- | --- |
| **EssTFNet** | 0.7644 | 0.7828 | 0.8336 | 0.8212 |
| DNA+Protein | 0.67307 | 0.6277 | 0.7263 | 0.7265 |
| DNA | 0.6298 | 0.6462 | 0.6803 | 0.6642 |
| No ATFNet | 0.7580 | 0.7512 | 0.8206 | 0.8022 |
| No LMs | 0.7467 | 0.7435 | 0.8028 | 0.7873 |
| Original NT v2 | 0.7612 | 0.7638 | 0.8230 | 0.8092 |

<sup>1</sup>DNA+Protein: Without ATFNet and physicochemical features; uses DNA and protein embeddings only. DNA: Uses DNA embeddings only. No ATFNet: Replaces ATFNet with a multi-head attention

When the model was restricted to use only DNA and protein features without the specialized architecture (denoted as DNA+Protein), a substantial decline in performance was observed, with accuracy dropping to 0.6731 and the F1-Score to 0.6277. The degradation became even more pronounced when the model relied solely on DNA features, resulting in the lowest recorded scores, including an accuracy of 0.6298 and an AUC of 0.6803. These findings emphasize the critical role of multi-modal integration in enhancing model effectiveness.

We further evaluated the impact of individual architectural components. Removing the ATFNet module (No ATFNet) led to significant declines in F1, AUC, and AUPR metrics, underscoring the critical role of the time-frequency network module in modeling long sequences. Similarly, the absence of language models (No LMs) also resulted in a noticeable drop in performance, highlighting the importance of pre-trained models in enhancing feature representation. While the original Nucleotide Transformer v2 (Original NT v2) demonstrated relatively strong performance, it still fell short compared to the full EssTFNet model. These experimental results collectively validate the necessity of each module within the architecture and emphasize their substantial contributions to boosting predictive performance.

### 2.5 Web server

To enhance the accessibility and usability of EssTFNet, we developed a fully-featured and user-friendly web server (https://esstfnet.art/). This platform integrates essential gene prediction models for 1,090 different human cell lines, based on EssTFNet, and aims to provide researchers with convenient, fast, and high-precision prediction services. For the benefit of the community and researchers, we have made our source code openly available through a public repository: https://github.com/QIANJINYDX/EssTFNet. The models and datasets are also accessible at: https://zenodo.org/records/15552651.

The workflow is illustrated in Figure 5. Users simply need to upload or enter the target DNA sequence and select the corresponding cell line from the dropdown menu to perform model predictions online. In cases where specific cell line information is unavailable, the platform also offers a general-purpose model suitable for preliminary analysis and exploratory studies across diverse experimental conditions. Upon submission, prediction results are automatically generated and presented in a dedicated results panel. In addition to the predicted gene scores, the server also integrates the DeepLIFT algorithm to assess the impact of each base pair on the model’s prediction results. This attribution analysis helps users understand which regions of the sequence are most critical to the model’s judgment, providing important references for further functional annotation and experimental design. The platform provides strong computational support for gene function research, target screening, and mechanism exploration, serving as an important bridge between artificial intelligence and functional genomics research.

**Figure 5.**
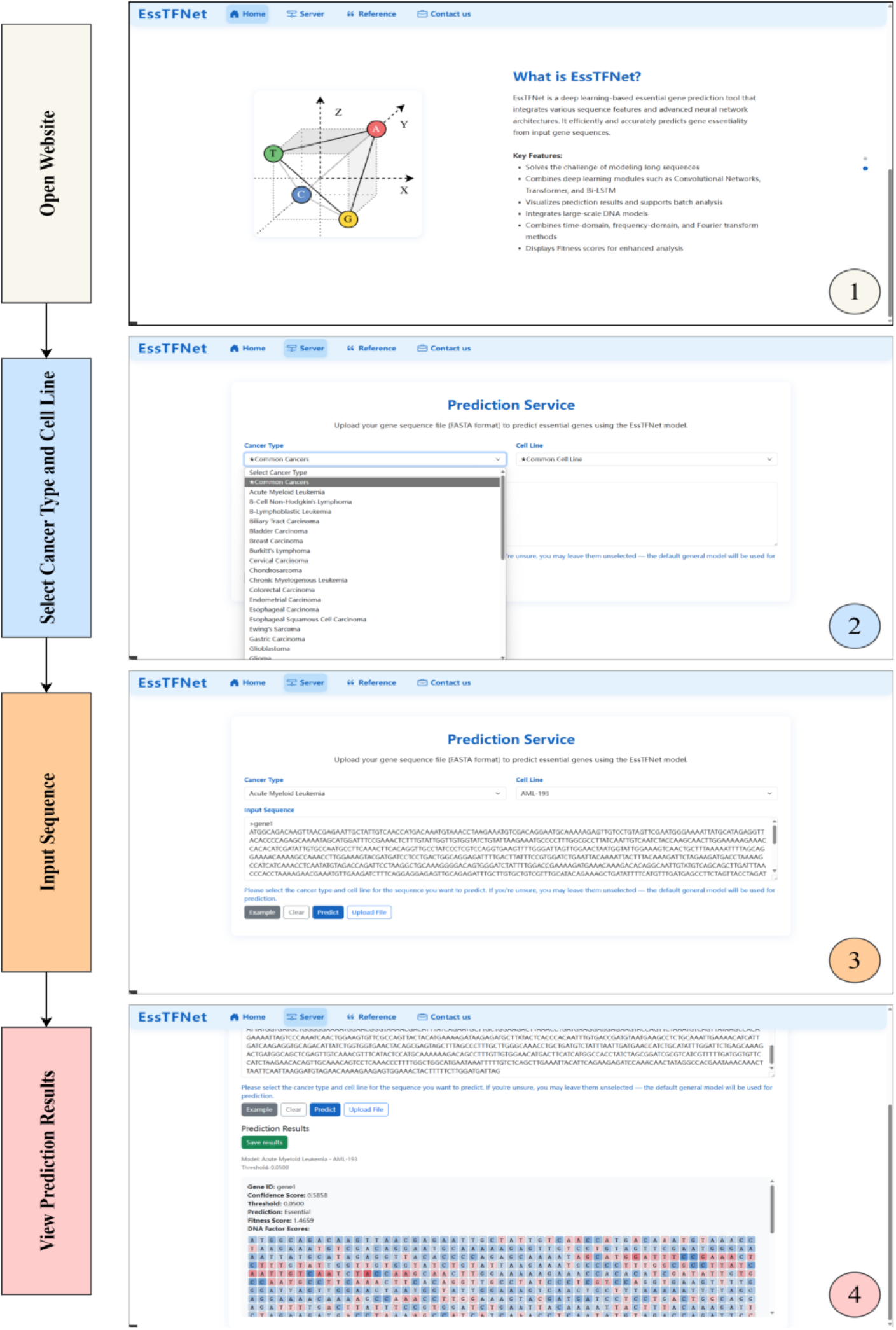
Step-by-Step Guide for Essential Gene Sequence Prediction

## 3 Discussion

Although experimental results demonstrate that EssTFNet achieves a good balance among prediction accuracy, model interpretability, and cross-species generalization capability—significantly outperforming current mainstream sequence-based deep learning methods—we must still acknowledge its limitations. The current framework primarily focuses on modeling and analyzing DNA sequences, and has made notable progress in mining DNA information through time-frequency mapping and integration with language models. However, in the process of feature attribution and mechanistic interpretation, EssTFNet does not fully consider sequence and structural features at the protein level. Important biological information such as conserved amino acid regions in translated proteins, secondary structures, and functional sites has yet to be systematically integrated.

Therefore, future work will focus on the following directions: (1) Cross-modal modeling: Introduce protein sequence and structural information to enable end-to-end modeling from transcription to translation by integrating DNA and protein language models. (2) Multi-omics data integration: Incorporate multi-omics data such as transcriptomics and epigenomics into model training to enhance predictive performance and generalization under specific biological contexts. (3) Interpretability mechanism optimization: Further incorporate graph neural networks (GNNs) and structural attribution methods to improve the model’s interpretability at the spatial structural level. (4) Expansion of task applicability: Explore the applicability of EssTFNet to other gene function prediction tasks, such as non-coding RNA function prediction (Ru et al. 2019) and cancer driver gene identification, to enhance its generality and practicality.

In summary, EssTFNet provides an efficient, interpretable, and extensible new framework for sequence-based functional gene identification. By integrating multimodal information and structural knowledge in the future, it holds great promise for advancing the application of deep learning methods in gene function modeling.

## 4 Materials and methods

### 4.1 Data collection and preprocessing

This study utilized a total of five datasets for the analysis. Detailed descriptions of datasets of interest are provided in Table 2.

**Table 2.** Summary of Essential Gene Datasets (S1–S5)

| Dataset | Data Content | Number of DNA Sequences | Coverage | Labels |
| --- | --- | --- | --- | --- |
| S1 | DNA and Protein | 11,899 | 11 cancer cell lines | Binary labels |
| S2 | DNA and Protein | 16,408 | 323 cancer cell lines | Binary labels |
| S3 | DNA and Protein | 16,335 | 1,090 cancer cell lines | Binary labels<br>Fitness scores |
| S4 | DNA and Protein | 16,335 | Same cell lines as S3 | Binary labels<br>Fitness scores |
| S5 | DNA | 478,431 | Cross-species essential gene data | Unsupervised (for MLM pre-training) |

The S1 dataset used in this study was originally constructed by Guo et al (Guo et al. 2017) in 2017, which integrates essential gene data from three independent studies covering 11 human cancer cell lines (A375, DLD1, GBM, HAP1, HCT116, HeLa, Jiyoye, KBM7, K562, Raji, and RPMI-8226) (Blomen et al. 2015; Hart et al. 2015; Wang et al. 2015). The study employed the majority rule method to define core essential genes, whereby a gene was labeled as essential if it was identified as such in more than six cell lines. Gene annotation information was retrieved from the HGNC database, and genes not included in the essential set were labeled as non-essential, forming a well-defined negative sample set (Seal et al. 2023).

The S2 and S3 datasets are derived from the Project Score initiative and the dataset released by Pacini et al. (Dwane et al. 2021; Pacini et al. 2021), respectively. The S2 dataset contains 16,408 gene sequences from 323 cell lines, while the S3 dataset contains 16,335 gene sequences from 1,090 cell lines. These two datasets provide rich and diverse gene expression information across various cell lines, which helps enhance the generalization ability of essential gene prediction models and their adaptability in different cellular environments.

The S4 dataset was constructed based on the S3 dataset, by applying the same majority rule method as in the S1 dataset to define core essential genes. In addition, the regression targets in S4 were defined as the average fitness scores across cancer cell lines in S3, providing a quantitative measure of gene essentiality. This resulted in a more representative subset of essential genes, which was used to build a generalizable essential gene prediction model.

The S5 dataset designed for building a DNA language model for essential genes was compiled by aggregating sequences from multiple essential gene databases, including DEG, CEG, Epath, and OGEE3 (Kong et al. 2019; Liu et al. 2020; Gurumayum et al. 2021; Luo et al. 2021). All sequences were preprocessed by converting them to uppercase and retaining only the four canonical nucleotides (A, G, C, and T), while removing spaces, newlines, and other non-standard characters. The dataset was then split into training and validation sets at a 0.95:0.05 ratio for pretraining on the masked language model (MLM) task.

To enhance the biological representation of gene data, we further integrated full-length protein sequence annotations for the S1, S2, and S3 datasets using the Consensus CDS (CCDS) database (Pujar, et al., 2018). Considering the extreme length distributions in the original DNA sequences (with some sequences exceeding 40,000 bp), the Isolation Forest algorithm (contamination ratio = 0.01) was applied to detect and remove exceptionally long sequences. After processing, 99% of the DNA sequences had lengths concentrated within 10,000 base pairs (as shown in the Figure 2C), significantly optimizing the input feature space for subsequent deep learning models. It is particularly noteworthy that the S3 dataset not only includes binary essential gene labels but also provides continuous fitness scores based on multi-cell line validation.

### 4.2 Model architecture

In this study, we designed a modular architecture, EssTFNet, to ensure the efficient extraction and integration of different feature information. The entire model consists of three main modules: the physicochemical feature extraction module, the sequence feature extraction module, and the pre-trained model integration module. The data flow begins with the preprocessing of DNA and protein sequences, where all sequences are standardized and converted into formats suitable for model input, including one-hot encoded sequences for both types (DNA and protein), and physicochemical feature vectors. The preprocessed data is then fed in parallel to each module for feature extraction. In the final output layer, the model concatenates the feature vectors from each module and uses a Sigmoid activation function to generate prediction values for essential genes, thereby completing the prediction process. The full architecture of the proposed EssTFNet model is schematically illustrated in Figure 6.

**Figure 6.**
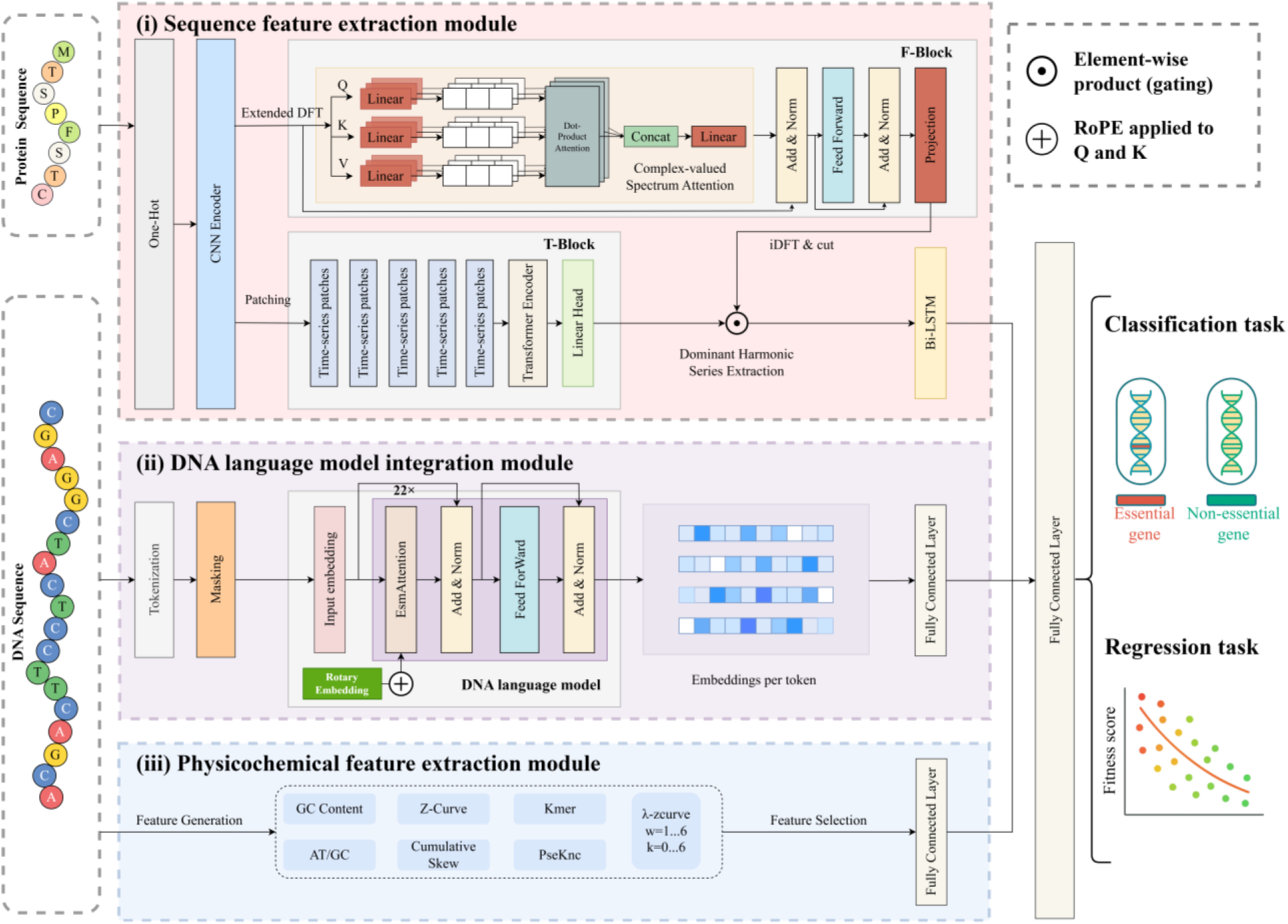
Framework of EssTFNet. EssTFNet has a hybrid architectural design, which efficiently integrates complementary representations from multiple branches. (i) Protein and DNA sequences are first encoded and processed by a CNN (TextCNN) to capture local motif patterns. The resulting features are further modeled by a T-Block for sequential dependencies and an F-Block that performs complex-valued spectrum attention in the time–frequency domain, followed by dominant harmonic series extraction and a Bi-LSTM module. (ii) In parallel, the DNA sequence is fed into a DNA language model (Nucleotide Transformer-v2) to obtain contextual token embeddings, followed by a fully connected layer. (iii) The physicochemical and sequence-derived features of DNA sequence (e.g., GC/AT content, k-mer, Z-curve, skew, PseKNC, etc.) are incorporated into a feature vector through a feature-selection module, followed by a fully connected layer. All feature representations are concatenated and fused through fully connected layers to produce multi-task outputs, including binary essential-gene classification (via a sigmoid activation) and continuous gene essentiality/fitness score regression.

#### 4.2.1 Time-frequency network design

When processing DNA and protein sequences, we treat the one-hot encoded sequences as time series and extract both their temporal and frequency information. For extracting this information, we adopted the methods from ATFNet (Ye et al. 2024) . For an input sequence *S* ∈ ℝ^*L*^ , the T-Block directly processes the sequence in the time domain, generating the output *S*_*o*_^(*t*)^ ∈ ℝ^*T*^. We use the extended Discrete Fourier Transform (DFT) to transform *S* into the frequency domain, generating the extended spectrum *F* ∈ ℂ^*L*+*T*^ . Then, through the Inverse Discrete Fourier Transform (iDFT), the spectrum is converted back to the time domain, and the F-Block generates the output *S*_*o*_^(*f*)^ ∈ ℝ^*T*^ . Finally, the outputs from the T-Block *S*_*o*_^(*t*)^ and the F-Block *S*_*o*_^(*f*)^ are combined through a weighted sum to generate the final output *Ŝ* ∈ ℝ^*T*^ . These weights are derived based on the energy of the fundamental harmonic sequence. The entire process can be formalized as:

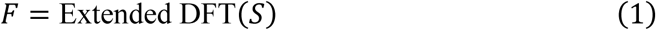

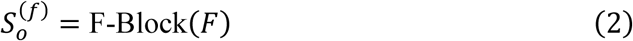

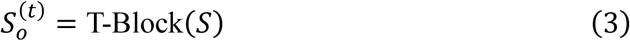

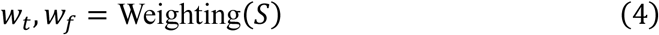

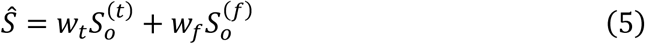

Fourier Transform (Extended DFT). It overcomes the limitations of input length, allowing us to obtain the input spectrum aligned with the DFT frequency bins of the full sequence. The specific formula is as follows:

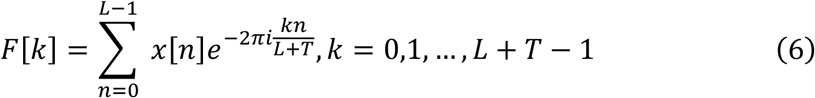

By using Equation 6, we obtain a spectrum of length L+T, aligned with the DFT spectrum of the full sequence. For real-valued time series, the conjugate symmetry of the output spectrum is an important feature of the DFT. Utilizing this property, computational cost can be reduced by considering only the first half of the output spectrum, as the second half provides redundant information.

The architecture of the F-Block is based on the Transformer encoder, with all parameters being complex-valued. Specifically, all computations within the F-Block are performed in the complex domain. The F-Block receives the uni-variate frequency spectrum 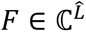 generated by the extended DFT as input, where 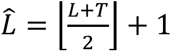 represents the length of the spectrum.

First, *F* is normalized by subtracting the mean and dividing by the standard deviation, with the standard deviation calculated from 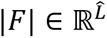. Subsequently, the mean is added back at the end of the structure. Next, 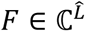 is embedded into *F*_*d*_ ∈ ℂ^*D*^ using a trainable linear projection 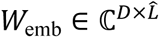. Since there is less temporal dependence within the frequency domain spectrum, the use of position embeddings is disabled.

Complex-Valued Spectrum Attention. An improved multi-head attention mechanism is used instead of the original attention mechanism. For each head *h* = 1,2, … , *H* , the embedded spectrum *F*_*d*_ ∈ ℂ^*D*^ is projected onto the frequency dimension *d* through a trainable projection. Specifically, 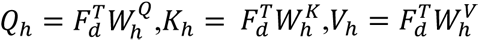, where 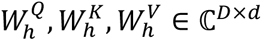 are learnable parameters. For each head, complex-valued dot-product attention is performed:

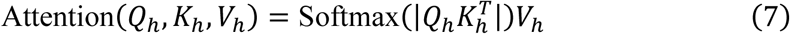

Then, the final output of the complex-valued spectrum attention is computed as follows:

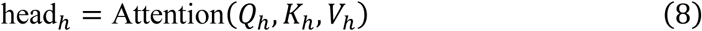

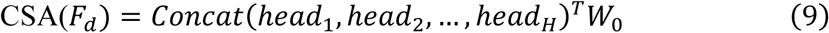

Where *W*_*O*_ ∈ ℂ^*hd*×*D*^ is a learnable parameter.

Following the design of LayerNorm and FeedForward layers in the Transformer, and extending to the complex domain in the residual connections, the output *F*_*z*_ after *M* layers of encoders is linearly projected to 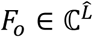. Then, we convert *F*_*o*_ back to the time domain using the inverse DFT (iDFT) and take the last *T* points (the prediction part) as the final output *X*_*O*_^(*f*)^ ∈ ℝ^*T*^ of the F-Block. The F-Block applies a full complex-valued neural network (CVNN) architecture (Barrachina et al. 2023).

The T-Block is responsible for capturing local dependencies in time series, which is more easily handled in the time domain. Chunking is an intuitive and effective method for capturing local dependencies in time series. This chunking is then combined with the Transformer encoder architecture to form the T-Block. First, it splits the input sequence *X* ∈ ℝ^*L*^ into multiple small chunks *X*_*p*_ ∈ ℝ^*P*×*N*^ , where *P* is the length of each chunk and *N* is the number of chunks. The small chunks *X*_*p*_ are then embedded and fed into the Transformer encoder. Through a linear projection, the final output *X*_*O*_^(*t*)^ ∈ ℝ^*T*^ is generated.

#### 4.2.2 Design of essential gene language model

We evaluated various existing DNA language models on the S4 general essential gene dataset, including DNABERT, DNABERT-2, GROVER, Nucleotide Transformer-v1, Nucleotide Transformer-v2 (NT v2), and Gena-LM-BERT (Ji et al. 2021; Zhou et al. 2023; Sanabria et al. 2024; Dalla-Torre et al. 2025; Fishman et al. 2025). Among these, NT v2 demonstrated the best performance in terms of the AUC metric and was therefore selected as the base model for further development, as shown in the results Figure 2E. Building on this, we further pre-trained the model using the S5 dataset to enhance its ability to recognize and understand essential genes. During the pre-training process, we set the model’s maximum supported sequence length to 2048 to ensure the capture of long-range contextual information. The vocabulary size was set to 4107, with input processing based on a 6-mer encoding scheme, which more effectively represents local structural features in biological sequences. The model underwent a total of 1,267,000 training steps, with training halted once the loss stabilized to prevent overfitting and ensure the model converged to an optimal state.

### 4.3 Extraction of physicochemical features from DNA sequences

In the process of physicochemical feature extraction, we selected a variety of features, including K-mer, PseKNC features, cumulative skew, Z-Curve, GC content, AT/GC ratio, and λ-Z-curve features, with the λ-Z-curve feature being validated as an effective feature in essential gene studies(Guo et al. 2017); see Table 3 for a detailed breakdown of feature types and quantities.

**Table 3.** Feature Types and Quantities.

| Feature Type | Number of Features |
| --- | --- |
| K-mer (k=1,2,3) | 84 |
| PseKNC (k=3) | 65 |
| Cumulative skew | 2 |
| Z-Curve | 3 |
| GC content | 1 |
| AT/GC ratio | 1 |
| λ-Z-curve (w=1...6, k=0...6) | 85,995 |
| Total | 86,151 |

To further optimize the feature set, we applied the RFE (Recursive Feature Elimination) method for feature selection on the S1 dataset. Considering the time overhead that large-scale feature selection might incur, we performed a step-by-step selection for the λ-Z-curve features. Specifically, we divided the 85,995 λ-Z-curve features into 9 groups, each containing 9,555 features, and combined them with other features like K-mer, so that the total number of features per group reached 9,711. Based on this, we selected the top 1,000 features from each group. After merging the 9 groups and removing duplicates, we obtained 8,263 features. We then applied the RFE method again to select the optimal features, and the results showed that when the number of features was set to 4,800, the model performance reached its best (as shown in Figure 2A). Therefore, we chose 4,800 features as the optimal number, providing efficient and streamlined input data for the model. Further analysis of these selected features revealed that the λ-Z-curve features demonstrated outstanding performance: among the 4,800 selected features, 4,730 were derived from the λ-Z-curve, indicating its dominant contribution to the final feature set (as shown in Figure 2B). This highlights the crucial role of the λ-Z-curve features in identifying and distinguishing essential genes, and suggests that they provide highly informative input that enhances the accuracy and efficiency of gene validation. For other datasets, we also used these 4,800 features as the input for the physicochemical features.

### 4.4 Model interpretability

To enhance the interpretability of our model and understand its internal feature representations and decision-making mechanisms, we employed two methods: t-distributed Stochastic Neighbor Embedding (t-SNE) (Maaten and Hinton 2008) for feature space visualization, and Deep Learning Important Features (DeepLIFT) (Shrikumar et al. 2017) for input attribution analysis. These approaches allow us to investigate how the model transforms and utilizes multi-level features to distinguish essential genes from non-essential ones.

#### 4.4.1 Feature representation visualization

To investigate the representational capabilities of our model across different feature levels, we employed t-distributed Stochastic Neighbor Embedding (t-SNE) for dimensionality reduction and clustering analysis. Specifically, we applied t-SNE to the output features of fully connected layers, extracted at various stages of the model. These included the physicochemical features derived from DNA sequences, the one-hot encoded features of protein sequences, and the high-level semantic representations from the final output layer.

t-SNE is particularly effective for visualizing high-dimensional data in a lower-dimensional space (typically two or three dimensions) while preserving local structures. It achieves this by minimizing the Kullback-Leibler (KL) divergence between two probability distributions: one representing pairwise similarities in the high-dimensional space and the other in the low-dimensional embedding. The objective function is given by:

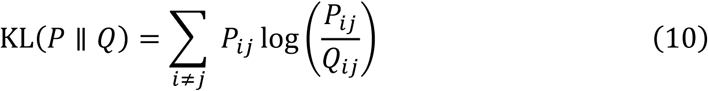

Where *P*_*ij*_ denotes the similarity between samples *i* and *j* in the original feature space, and *Q*_*ij*_ denotes the corresponding similarity in the embedded space.

By comparing the t-SNE clustering results across different feature representations, we aim to reveal how effectively the model separates different classes or patterns at various stages of processing. This analysis provides valuable insights into the model’s internal feature learning mechanisms and supports further interpretability of its decision-making process.

#### 4.4.2 DeepLIFT-based attribution analysis

To explain the model’s prediction decisions for a given DNA sequence based on specific features, we employed the DeepLIFT attribution method (Shrikumar et al. 2017). Processing procedure is illustrated in Figure 7. First, the DNA sequence is standardized to a fixed length of 1800 bp (which corresponds to the average sequence length range), with any insufficient parts filled with “N” characters. Next, based on the background base probability distribution of AGCT in the dataset, we construct the corresponding one-hot form baseline input. The constructed baseline is then inputted into the model along with the actual input, and DeepLIFT is used to calculate the attribution contribution of each position to the prediction result. Without altering the original model structure, this method allows for a refined evaluation of the importance of feature inputs.

**Figure 7.**
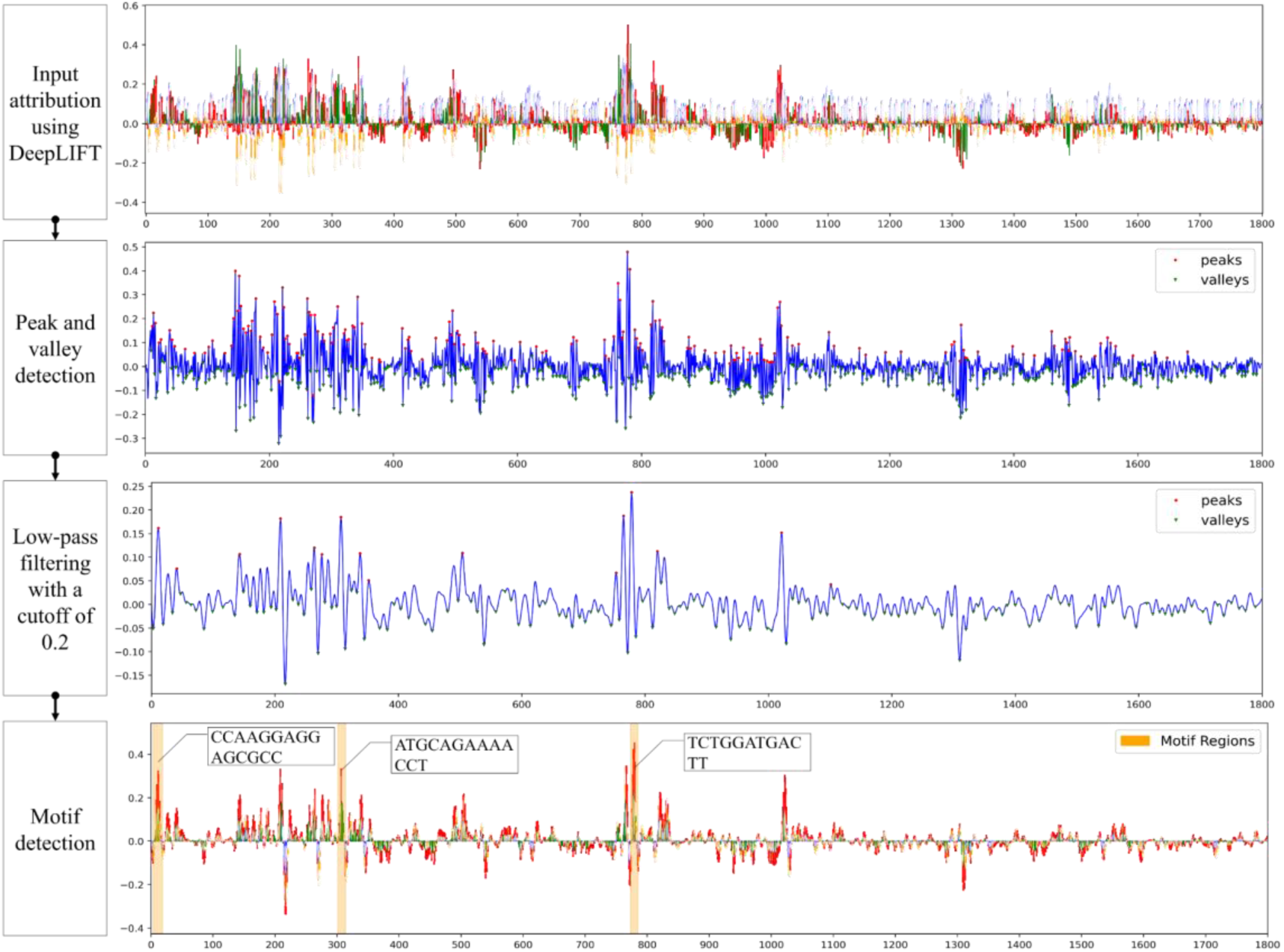
DeepLIFT attribution processing procedure. First, the DeepLIFT algorithm is used for feature attribution on the input sequence, visualizing the DNA sequence feature distribution with a colored fluctuating curve (with the x-axis representing the sequence and the y-axis representing the attribution strength). Then, key signal nodes are located based on extremum detection (with red/green dots marking peaks and valleys), and the main trend is extracted after smoothing with a low-pass filter at a cutoff frequency of 0.2. Finally, specific binding regions (such as sequences like CCAAGGAGGAGG) are located at the gene sequence level, highlighted by orange annotations.

It was found that longer motifs are crucial in human essential gene prediction tasks, as these long motifs often correspond to important sequence-specific regulatory elements, whose structural features are significantly correlated with gene functional necessity (Xie et al. 2007). Therefore, we integrated digital signal processing techniques into the motif extraction algorithm: by applying a tunable low-pass filter for frequency domain analysis of the sequences, and combining a peak detection algorithm, we achieved precise control over the motif length range.

The backpropagation contribution scores, after signal preprocessing, are fed into a low-pass filter. Unlike a high-pass filter, which emphasizes local mutation features, the low-pass filter systematically suppresses high-frequency noise components through its amplitude-frequency characteristics, while retaining the basic low-frequency features that represent important regulatory patterns. This enables the stable extraction of continuous feature intervals with significant biological meaning from the complex contribution scores. Below is the transfer function of a first-order analog low-pass filter:

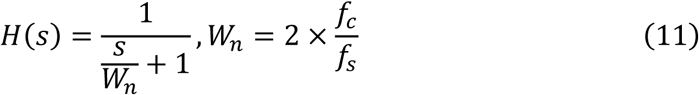

In this case, *f*_*c*_ is the cutoff frequency, and *f*_*s*_ is the sampling frequency. The study uses an eighth-order Butterworth low-pass filter to enhance frequency-domain selectivity. The normalized cutoff frequency parameter *W*_*n*_ is dynamically configured according to the target motif length: for detecting motifs of lengths 30 bp, 10 bp, and 5 bp, the corresponding threshold values are 0.1, 0.2, and 0.4, respectively. This filter effectively extracts long motif features rich in high-frequency components by suppressing short-term fluctuations (high-frequency noise) while retaining the long-term trend signals dominated by low frequencies.

### 4.5 Motif screening

To identify sequence motifs significantly associated with essential genes, we performed motif enrichment analysis within a hypergeometric hypothesis testing framework. For each candidate motif, Fisher’s exact test (one-tailed, directionality set for positive enrichment) was applied to a 2×2 contingency table (containing motif occurrence counts in positive/negative sample groups and total sequence numbers per group) to assess whether its frequency in the positive sample group significantly exceeded that in the negative group. Raw p-values were subsequently adjusted for false discovery rate (FDR) using the Benjamini-Hochberg procedure to mitigate multiple hypothesis testing artifacts (Thissen et al. 2002). To enhance biological relevance, effect size screening criteria were implemented: odds ratios (OR) with 95% confidence intervals (CI) were calculated, retaining only motifs meeting the thresholds of adjusted p-value <0.05, OR >2, and CI lower bound >1 as significantly enriched. This dual-filter strategy, integrating statistical significance and effect magnitude thresholds, ensures the identified motifs exhibit both statistical robustness and functional correlation. For detailed results, please refer to Section 3.2 of the Results.

### 4.6 Implementation details

This study uses the holdout method to validate the model on the benchmark dataset. To address the limitations of traditional evaluation methods, we improved the data splitting strategy. In previous experiments, researchers commonly used stratified sampling to split the training and testing sets based on the ratio of positive and negative samples. However, the essential gene dataset in this study suffers from a class imbalance issue (with significantly more non-essential gene samples than essential genes). If the traditional stratified splitting method were used, the testing set would also have an imbalanced distribution, making it difficult to accurately assess the prediction performance for essential genes. Therefore, we constructed the test set through balanced sampling - randomly selecting 20% of the essential gene samples and matching them with an equal number of non-essential gene samples to form an independent test set. The remaining samples were all included in the training set. We used five-fold cross-validation for model training, with Adam as the optimizer (Adam 2014). For classification tasks, we used binary cross-entropy (BCE), while for regression tasks, mean squared error (MSE) was applied.

All models were implemented using the Pytorch and Scikit-learn libraries. Training was conducted on a workstation equipped with an Intel(R) Xeon(R) Silver 4310 CPU @ 2.10GHz with 256GB RAM and 4 NVIDIA GeForce RTX 4090D GPUs.

### 4.7 Evaluation metrics

This study uses multiple evaluation metrics to comprehensively assess model performance. For classification tasks, ACC (accuracy) is used to evaluate overall prediction accuracy, the F1 score is used to address class imbalance by considering both precision and recall, and AUC (Area Under the ROC Curve) and AUPR (Area Under the PR Curve) are used to evaluate the model’s classification ability at different thresholds and its performance on minority classes(Guo et al. 2024; Huang et al. 2024; Huang et al. 2025). For regression tasks, the coefficient of determination (R²) is used to measure the model’s goodness of fit, while the root mean square error (RMSE) quantifies the deviation between predicted and true values. These complementary metrics ensure the comprehensiveness and reliability of the evaluation results from different dimensions.

## Acknowledgments

This research was funded by the National Natural Science Foundation of China (Grant No. 62502073, 62172078) and the Fundamental Research Funds for the Central Universities (No. ZYGX2024Z011).

## Author Contributions

D-X.Y. conceived and designed the study, developed the EssTFNet framework, and performed the experiments and analyses. S-S.Y., W.S., H-Q.Z., and R.L. contributed to data preprocessing, physicochemical feature engineering, and experimental evaluation. D-X.Y. and R.L. carried out model benchmarking and statistical analyses. W.S. and Y-C.Q. contributed to software implementation and web server deployment. N.D., H.L., and Y-T.J. supervised the project and provided critical revisions to the manuscript. D.Y. drafted the manuscript with input from all co-authors. All authors read and approved the final manuscript.

## Institutional Review Board Statement

Not applicable.

## Informed Consent Statement

Not applicable.

## Data Availability Statement

The source code openly available through a public repository, https://github.com/QIANJINYDX/EssTFNet.

